# BRIDGE: Biological Antimicrobial Resistance Inference via Domain-Knowledge Graph Embeddings

**DOI:** 10.64898/2026.02.09.704676

**Authors:** Aravind Iyer, Yusuff Kazeem, Enayat Rajabi, Somayeh Kafaie

## Abstract

Antimicrobial resistance (AMR) is a growing global health crisis, responsible for an estimated 1.27 million deaths in 2019 alone. Traditional approaches to identifying antibiotic resistance genes (ARGs) are often labour-intensive and limited in their ability to detect novel resistance mechanisms. In this study, we propose BRIDGE, a knowledge graph-based framework, to improve AMR gene prediction by integrating gene neighbourhood information and protein-protein interaction (PPI) networks. Focusing on *Klebsiella pneumoniae* and *Escherichia coli*, we construct a comprehensive and biologically grounded knowledge graph using curated data from CARD, STRING, and DrugBank. We apply knowledge graph embedding models which are fed into neural networks to infer novel AMR links, achieving classification accuracy of up to 97%. Our results demonstrate that incorporating biologically meaningful relationships, such as gene neighbourhood information and PPIs, enhances the predictive accuracy and interpretability of AMR link predictions. This work contributes to the development of scalable and data-integrated approaches for advancing antimicrobial resistance surveillance and drug discovery. BRIDGE implementation and data are available at https://github.com/GraphML-lab/BRIDGE.

## I. Introduction

**A**NTIMICROBIAL Resistance (AMR) has become a burden on healthcare practitioners because it threatens the use of antibiotics to eliminate bacterial infections in patients [1]. In Canada in 2018, it was estimated that AMR was directly responsible for around 5,400 deaths, and by 2050, if the resistance rates stay at 26% or increase to 40%, AMR could cause 7,000-13,700 Canadian deaths [2]. AMR occurs when the infection-causing microorganisms (e.g., bacteria) mutate or receive antimicrobial resistance genes (ARGs) through horizontal gene transfer (HGT) [3]. HGT involves the transfer of genes without inheritance from parents to offspring. Detecting ARGs and HGT cases is a critical step in developing strategies to combat AMR and improving the efficacy of antibiotics. The traditional methods for identifying ARGs are resource-intensive and time-consuming. They are limited in detecting new resistance patterns since these methods depend on prior knowledge of the mutations that confer resistance.

Recently, there has been an emergence of deep learning-based and bioinformatic approaches, including the use of knowledge graphs (KGs), that overcome the limitations of traditional methods for identifying ARGs. In particular, the Knowledge Integration and Decision Support (KIDS) frame-work [4] implemented a KG to identify new relationships between *E. coli*-associated genes and antibiotics.

In addition, Gao et al. [5] used electronic health records and graph-based modelling to generate patient-specific AMR treatment recommendations, while Quemeneur et al. [6] introduced a multi-modal, temporal KG aligned for resistance surveillance. Furthermore, Islam et al. [7] proposed a comprehensive KG-based framework to detect and analyse HGT events contributing to AMR. These studies highlight the growing potential of KGs for integrating clinical and environmental data into AMR prediction workflows.

Despite these advances, existing KG frameworks often suffer from limitations in capturing biological context or strain-level specificity. For example, models such as GNN-ARG [8] and MetagenomicKG [9] focus on metagenomic and gene prediction aspects but do not tightly link structural gene neighbourhoods with curated interaction networks.

In this study, we propose BRIDGE, a more comprehensive and controlled AMR prediction framework, by constructing a KG that incorporates AMR gene neighbourhoods and protein-protein interaction (PPI) networks. Unlike existing methods that require inconsistency resolution due to heterogeneous data sources, our approach ensures data reliability by carefully curating the KG from verified sources.

Our primary contributions include (1) the development of a structured KG that integrates AMR gene neighbourhoods, drug-target relationships, and PPIs among other relationships; (2) an improved prediction model for limited-data AMR genes using KG embeddings and neural networks; and (3) a comparative evaluation of our method against existing approaches to demonstrate its effectiveness in predicting novel AMR associations.

The remainder of this paper is organised as follows. Section 2 provides an overview of related work in AMR detection. Section 3 details our methodology, including data sources, KG construction, and link prediction methods. Section 4 presents experimental results and comparative evaluations. Finally, Section 5 discusses our findings, potential limitations, and future research directions.

## II. Background and Related Work

Traditional approaches for identifying AMR rely on culture-based methods and genome sequencing, but they are time-consuming, expensive, and often fail to predict emerging resistance patterns [10]. KGs have gained significant attention in biomedical research, as they offer a structured representation of entities and their relationships. Each relationship is a (subject, predicate, object) triplet linking two entities. Several studies have explored the use of KGs for biological network analysis, especially for drug combination therapies [11], metagenomic analysis [9], drug discovery and drug repurposing [12]–[16], suggesting that these methods could be further refined for AMR research. For example, Ye et al. [11] proposed a multi-modal deep learning framework integrated with a KG to predict drug synergy for infectious diseases, and PHPGAT [17] used attention-based GNNs on heterogeneous AMR KGs to predict phage-host associations.

In the context of AMR, KGs can integrate diverse data types, such as drug interactions, PPIs, and ARGs, enabling the discovery of novel relationships that might not be evident through traditional analysis. A notable example is the Knowledge Integration and Decision Support (KIDS) frame-work [4], which constructs a KG by aggregating data from multiple sources and resolving inconsistencies through automated reasoning. This framework, which has been applied to the biological domain of *E. coli*, integrates 10 different sources of data. These sources form an intermediate KG that is refined using an inconsistency resolver. The inconsistencies are introduced as a result of including multiple heterogeneous data sources and by assuming the inclusion of corrupted triplets. This approach demonstrated the potential of KGs in predicting unknown ARGs; however, its precision and F1-score remained suboptimal due to the challenges associated with integrating heterogeneous data and negative triplets.

While frameworks such as KIDS have demonstrated the utility of KGs, the limitations imposed by post-hoc inconsistency resolution due to the excessive triplet corruption and the lack of genomic context integration contribute to suboptimal performance. This gap in research is addressed by BRIDGE, our study that aims to enhance knowledge graph-based AMR prediction by constructing a more comprehensive and refined KG that incorporates AMR gene neighbourhoods and PPIs. The effectiveness of this approach is demonstrated by focusing on antibiotic resistance in *K. pneumoniae* and *E. coli. K. pneumoniae* is a significant species of bacterium causing a wide range of diseases such as cystitis, pneumonia, and endocarditis and is increasingly developing resistance to antibiotics [1]. *E. coli* is a gram-negative bacterium that is the predominant cause of urinary tract infections (approximately 70% of cases) and a major cause of sepsis worldwide. Many *E. coli* strains now harbour multidrug resistance mechanisms, making antibiotic-resistant *E. coli* an emerging global threat [18]. *E. coli* was chosen because of a large amount of data being present for the bacterial species across datasets.

By carefully curating data from validated sources such as STRING [19], CARD [20], and DrugBank [21], this study aims to eliminate the need for postprocessing inconsistency resolution, thereby improving the reliability and predictive accuracy of the KG. Unlike prior studies that integrate multiple inconsistent sources and require complex post-hoc reconciliation, our framework emphasises upfront biological consistency and contextual AMR gene neighbourhoods for deeper and more explainable predictions. Our approach extends previous efforts by incorporating an additional biological context, ensuring that predicted AMR relationships are not only computationally inferred but also biologically meaningful.

## III. Methodology

This section outlines the methods and procedures used in our study, including data collection, KG construction, and embedding techniques for link prediction. Figure 1 shows the flowchart that represents the entire process of this study from data collection to evaluation.

**Fig. 1.**
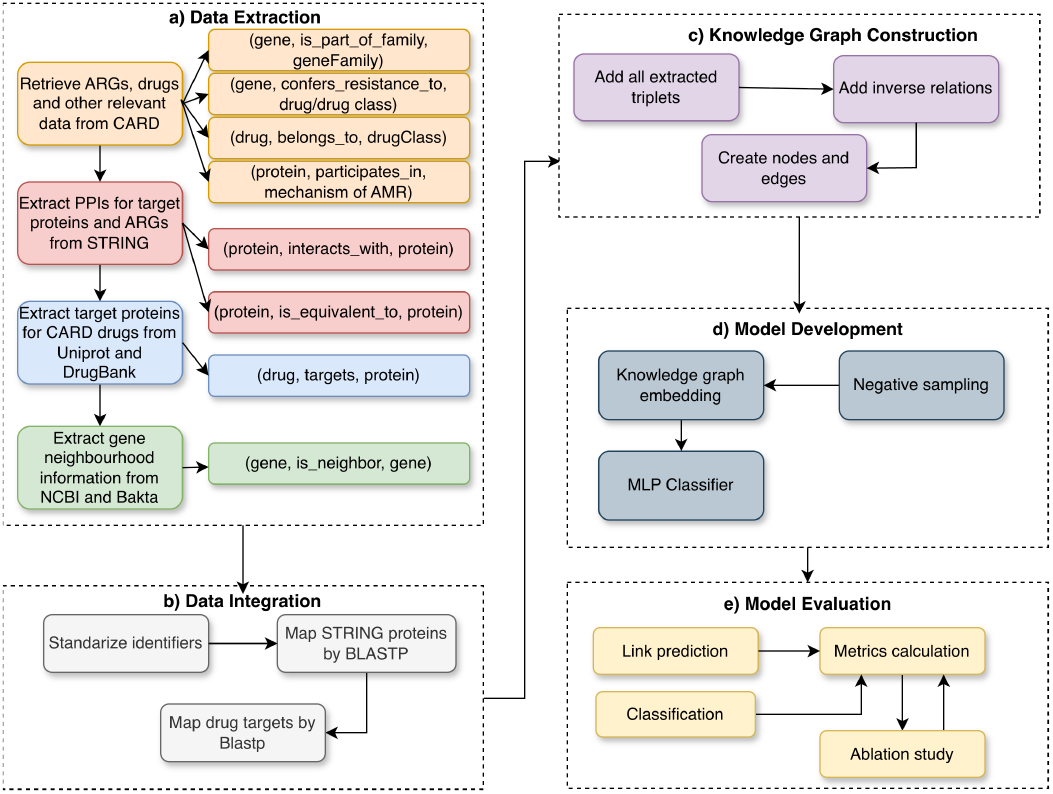
A flowchart representing the full pipeline of BRIDGE.

### A. Data Collection and Integration

To construct a comprehensive KG, we gathered data from multiple sources, ensuring a unified and consistent representation despite the heterogeneous nature of the datasets. Most of the data used in this study were pre-annotated, allowing us to focus on defining key relationships and integrating them into the KG. Data quality assurance was a priority, and all sources were thoroughly reviewed before integration.

The focus of our methodology is on ARGs and their associated metadata, including gene sequences, PPIs, and drugtarget relationships. The Comprehensive Antibiotic Resistance Database (CARD) served as the primary source for AMR-related information, while additional relationships were extracted from STRING, UniProtKB, and DrugBank. A summary of the data sources used in this study is provided in Table I, and the key relationships included in the KG are detailed in Table II.

**TABLE I.**
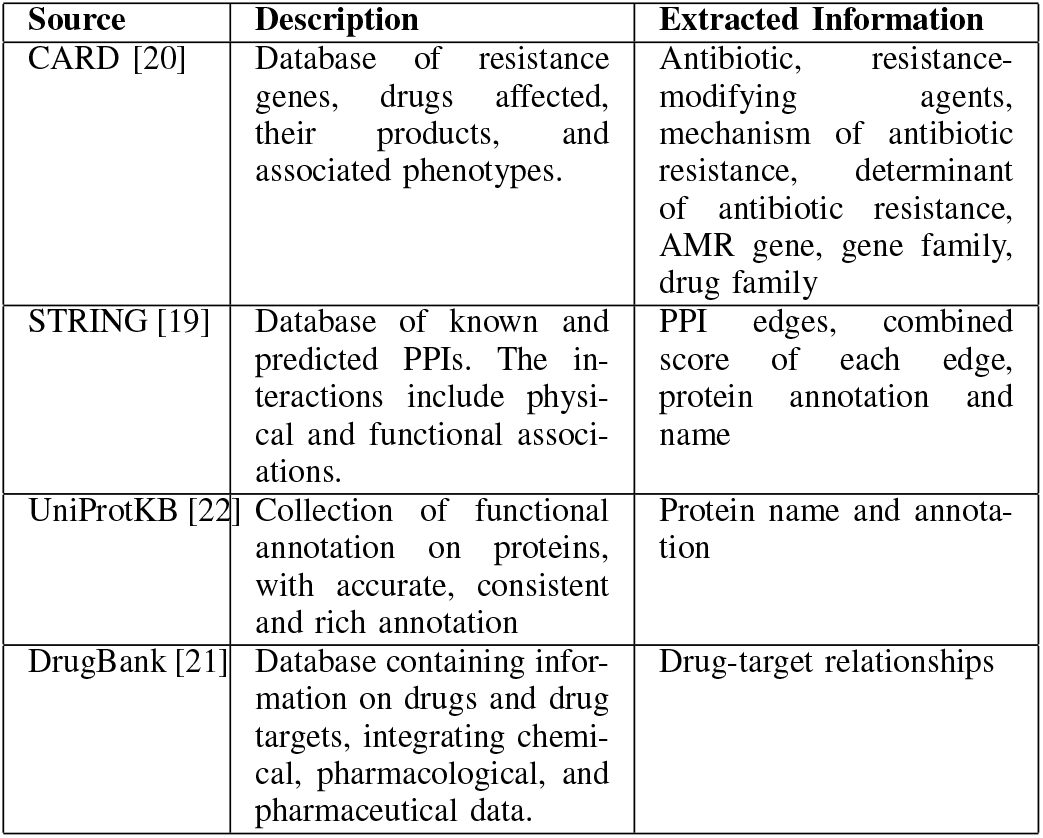
Data sources used for the proposed framework.

| **Source** | **Description** | **Extracted Information** |
|---|---|---|
| CARD [20] | Database of resistance genes, drugs affected, their products, and associated phenotypes. | Antibiotic, resistance-modifying agents, mechanism of antibiotic resistance, determinant of antibiotic resistance, AMR gene, gene family, drug family |
| STRING [19] | Database of known and predicted PPIs. The in-teractions include physi-cal and functional associ-ations. | PPI edges, combined score of each edge, protein annotation and name |
| UniProtKB [22] | Collection of functional annotation on proteins, with accurate, consistent and rich annotation | Protein name and annota-tion |
| DrugBank [21] | Database containing infor-mation on drugs and drug targets, integrating chemi-cal, pharmacological, and pharmaceutical data. | Drug-target relationships |

**TABLE II.**
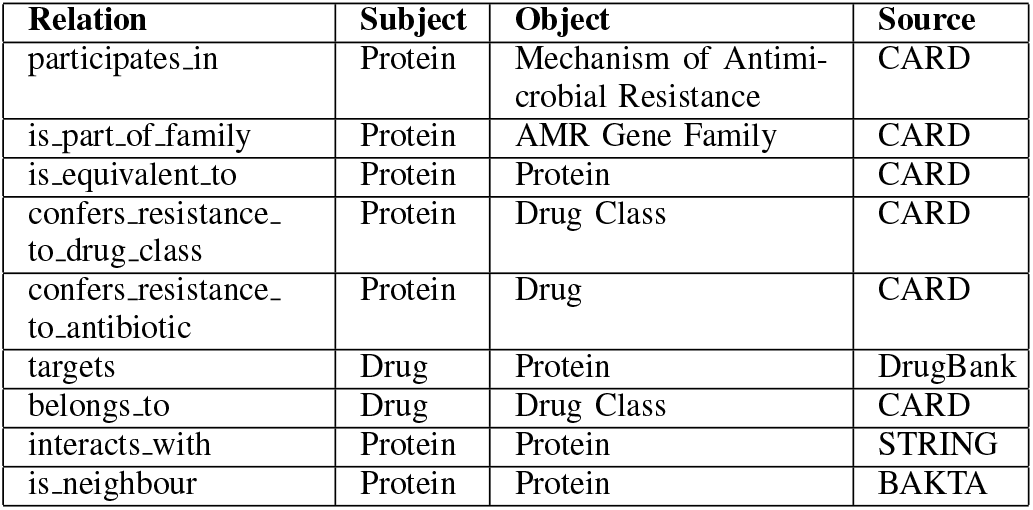
Entities and relations defined in the KG.

| **Relation** | **Subject** | **Object** | **Source** |
|---|---|---|---|
| participates_in | Protein | Mechanism of Antimi-crobial Resistance | CARD |
| is_part_of_family | Protein | AMR Gene Family | CARD |
| is_equivalent_to | Protein | Protein | CARD |
| confers_resistance_to_drug_class | Protein | Drug Class | CARD |
| confers_resistance_to_antibiotic | Protein | Drug | CARD |
| targets | Drug | Protein | DrugBank |
| belongs_to | Drug | Drug Class | CARD |
| interacts_with | Protein | Protein | STRING |
| is_neighbour | Protein | Protein | BAKTA |

Across all datasets, standard identifiers were mapped to common names to facilitate visual interpretation and down-stream analysis.

- **Core AMR Relationships:** Fundamental resistance relationships (e.g., participates in, confers resistance to) were retrieved directly from CARD using keyword-based searches.
- **Protein-Protein Interactions (PPIs):** PPIs reveal physical and functional cellular connections [23], which help identify novel resistance candidates [24]. To minimize noise while including meaningful interactions, we applied a threshold of 0.4 for the confidence score [25]. Because sequence representations vary, STRING proteins were mapped to CARD proteins using BLASTp with a 0.4 identity threshold [26]. STRING proteins lacking connections beyond *interacts with* were removed to prevent them from dominating the network.
- **Drug-Target Relationships:** Resistance frequently arises from modifications to drug targets [27], [28]. We mapped CARD-identified resistance drugs to their protein targets in DrugBank. Using UniProt sequences, we performed a BLASTp alignment (0.4 identity threshold [29]) against the existing CARD/STRING proteins in the KG, retaining only the top match per drug to form the final drug-protein pairs. The integration of drug-target relationships is shown in Figure 2.
- **Gene Neighbourhoods:** ARGs often cluster with uncharacterised genes to facilitate co-regulation and horizontal transfer [30]–[34]. To capture this context, we used BAKTA [35] to annotate ten NCBI genomic sequences per species. The annotations were processed using the Resistance Gene Identifier (RGI) [20], allowing us to extract the immediate genetic neighbors of known ARGs and map them to the KG via common name matching.

**Fig. 2.**
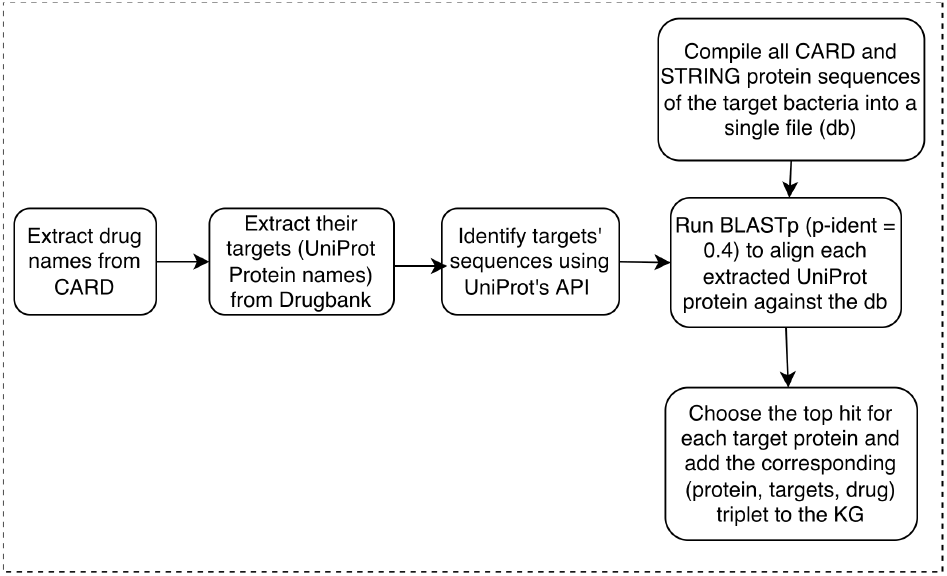
The steps taken to extract drug-target relationships.

### B. KG Construction

The KIDS’s KG served as a useful reference, and we adopted and enhanced parts of its schema by incorporating PPIs, gene neighbourhood information, and drug-target relationships. These enhancements provide the model with more relevant information, improving its ability to predict new links. The structure of the proposed KG is shown in Figure 3. Relationships derived directly from CARD are shown in red, while those extracted from external sources like DrugBank and STRING are shown in green, and the remaining relationships are depicted in black.

**Fig. 3.**
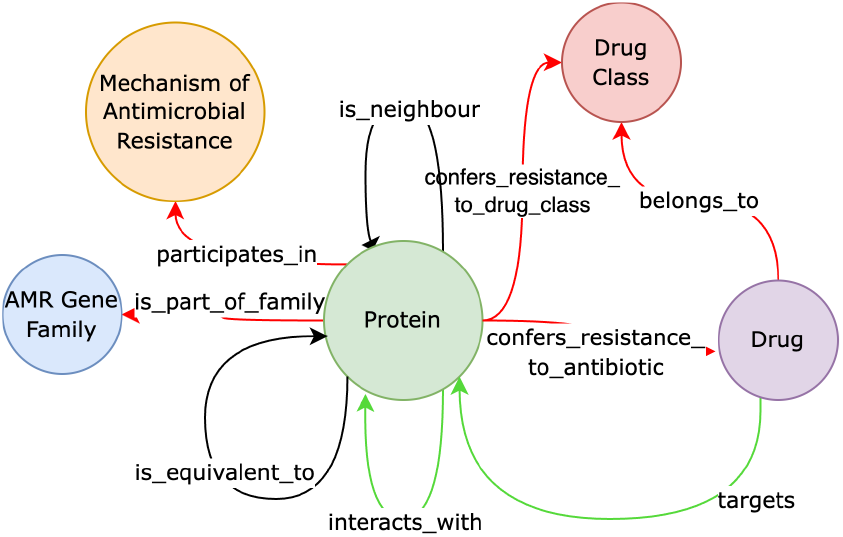
Underlying ontology structure for the KG.

### C. ARG Prediction

We employed KG embedding (KGE) models to learn vector representations of entities and relations and capture multi-relational patterns in the data. Specifically, we used both distance-based models (TransE [36], RotatE [37]) and semantic matching models (ComplEx [38], HolE [39], DistMult [40]) to leverage their complementary strengths in modeling relational structure while maintaining robustness on relatively small KG datasets. We explicitly incorporated inverse relationship patterns into the KG to ensure correct bidirectional reasoning, and used ComplEx to effectively model symmetric relations.

Many classical KGE models rely on linear or bilinear scoring functions and lack explicit non-linear transformations. To address this limitation, we attach a multilayer perceptron (MLP) to the learnt embeddings, enabling the model to capture non-linear patterns. In our setting, the MLP also reformulates the task as a binary classification problem by estimating the probability that a given protein–drug pair exhibits the *confers resistance to antibiotic* relationship, rather than producing a purely regression-based score.

The input to the MLP consists of three vector embeddings corresponding to the subject, relation, and object of each triplet. These embeddings are obtained from the preceding KGE model and concatenated to form a single input vector. To capture a compact representation of the triplet, this concatenated vector is first passed through a linear layer that compresses the input, effectively summarising the subject, relation, and object embeddings. The compressed rep-resentation is then combined with the original concatenated features and fed into two hidden layers. Each hidden layer employs ReLU activation, batch normalisation, and dropout to enhance model expressiveness and mitigate overfitting. The final output layer produces a single probability value, indicating whether the given triplet, specifically representing the *confers resistance to antibiotic* relationship between a protein and a drug, is predicted to be valid.

## IV. Evaluation and Results

### A. Experimental Setup

After data collection and integration, and constructing the KGs, the graph for *K. pneumoniae* consists of 1090 nodes with 9 unique relationships and 7403 edges. The *E. coli* KG consists of 812 nodes and 9267 edges.

We evaluated the link prediction capabilities of the constructed KG using multiple KGE models. The embeddings were generated using the AmpliGraph [41] library, and the models were trained on one NVIDIA A100 GPU (40 GB of memory) and two AMD EPYC 7413 CPU cores. For hyperparameter optimisation, we used AmpliGraph’s built-in grid search procedure to evaluate 20 different parameter combinations and identify the most effective configuration [42]. To ensure model stability and convergence, the number of training epochs was capped at 300 along with early stopping. This tuning process was performed independently for each KGE model to optimise their performance under distinct configurations. A dedicated validation set was used for each model to guide the selection of hyperparameters and prevent overfitting.

In addition to hyperparameter optimisation, *k*-fold cross-validation was employed. This approach partitions the dataset into *k* equal folds; in each iteration, one fold is used as the validation set while the remaining *k − 1* folds are used for training. Owing to the limited number of candidate test triplets for *K. pneumoniae* (351), *k* was set to 3 to ensure sufficient statistical reliability of the test results. In contrast, *E. coli*, with 896 candidate test triplets, allowed the use of *k* = 5. This strategy provides more robust performance estimates by reducing sensitivity to any single data split and supports more effective hyperparameter tuning by mitigating overfitting and promoting better model generalisation.

### B. Negative Sampling

Negative sampling was used to generate artificial invalid triples during training, enabling the KGE models to distinguish valid from invalid relationships and avoid overfitting to only positive examples. We adopted uniform negative sampling, where the head or tail of each positive triple is randomly replaced to create negative samples (ensuring that the generated negatives do not already exist in the dataset), as it is a simple, scalable, and widely used strategy that performs effectively across different datasets and KGE models.

### C. Evaluation Metrics

The performance of each KGE model was assessed using standard metrics, including Mean Rank (MR), Mean Recip-rocal Rank (MRR) and Hits@k. The classification tasks were evaluated using precision, recall, accuracy, F1-score, AUROC and AUPRC scores.

### D. Link Prediction

Table III summarises the performance of our framework using five different KGE models on KGs constructed for *K. pneumoniae* and *E. coli*. The results show that semanticmatching models (ComplEx and DistMult) consistently outper-form the distance-based models across all evaluation metrics. This suggests that semantic similarity modelling is particularly effective in capturing complex biological relationships within the AMR KG.

**TABLE III.**
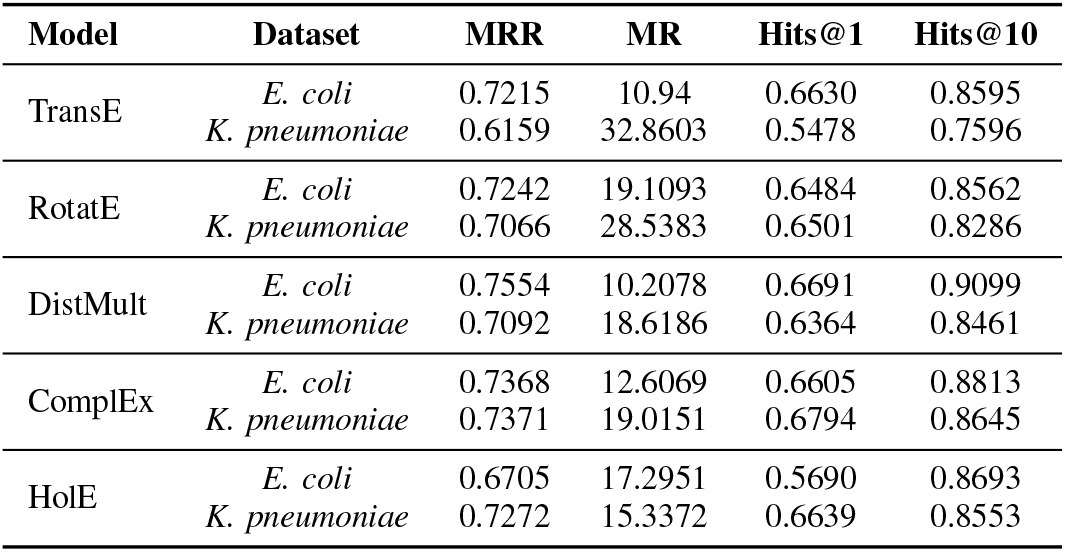
Performance of KGES on the *K. pneumoniae* and *E. coli* KGS.

To further enhance predictive performance, we augment each KGE model with a four-layer MLP, consisting of an input layer, two hidden layers, and an output layer. The MLP introduces non-linearity that enables more expressive modelling of complex biological interactions. It takes as input the concatenated subject, relation, and object embeddings, here focusing on the *confers resistance to antibiotic* relationship between proteins and drugs, and outputs a probability indicating whether a given triplet is valid. This formulation converts the link prediction task into a binary classification problem, allowing direct comparison with the KIDS framework [4] while encouraging generalisation beyond what standalone KGE models can achieve.

As shown in Figure 4, we compared BRIDGE against KIDS, a state-of-the-art method for AMR link prediction that is specifically designed for *E. coli*. Since KIDS is not available for other organisms, this comparison is restricted to *coli*, and the KIDS performance values are reported directly from the original manuscript. Under this setting, BRIDGE consistently achieves higher precision, recall, and F1 scores across multiple KGE backbones, with particularly notable improvements in precision, suggesting a reduced rate of false positive predictions when identifying AMR–drug associations. Figure 5 demonstrates that BRIDGE’s performance remains robust across different KGE models when applied to the *K. pneumoniae* KG. While some variation across embedding models is observed, particularly in recall, the overall performance remains consistently high, suggesting that the MLP can leverage diverse embedding structures to capture relevant biological signals. Additionally, Figure 6 summarises BRIDGE’s classification performance in terms of accuracy, AUROC, and AUPRC for both organisms. The consistently strong AUROC and AUPRC values support BRIDGE’s effectiveness as a link prediction framework.

**Fig. 4.**
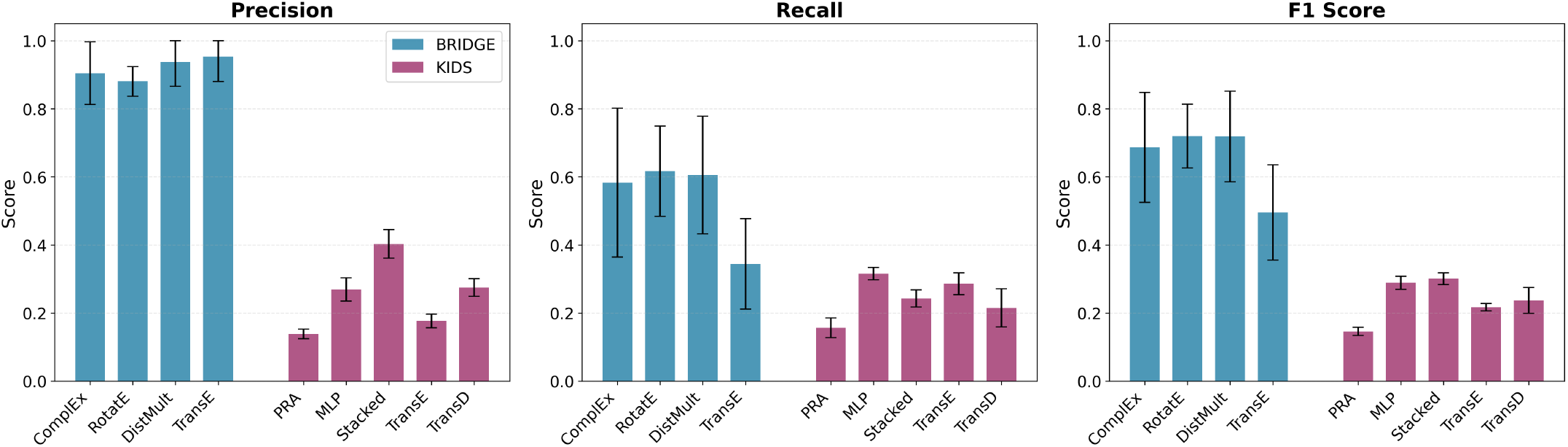
BRIDGE performance in *E. coli*: Comparison of link prediction results for BRIDGE vs. KIDS. BRIDGE utilizes an MLP model with inputs generated by several KGE models.

**Fig. 5.**
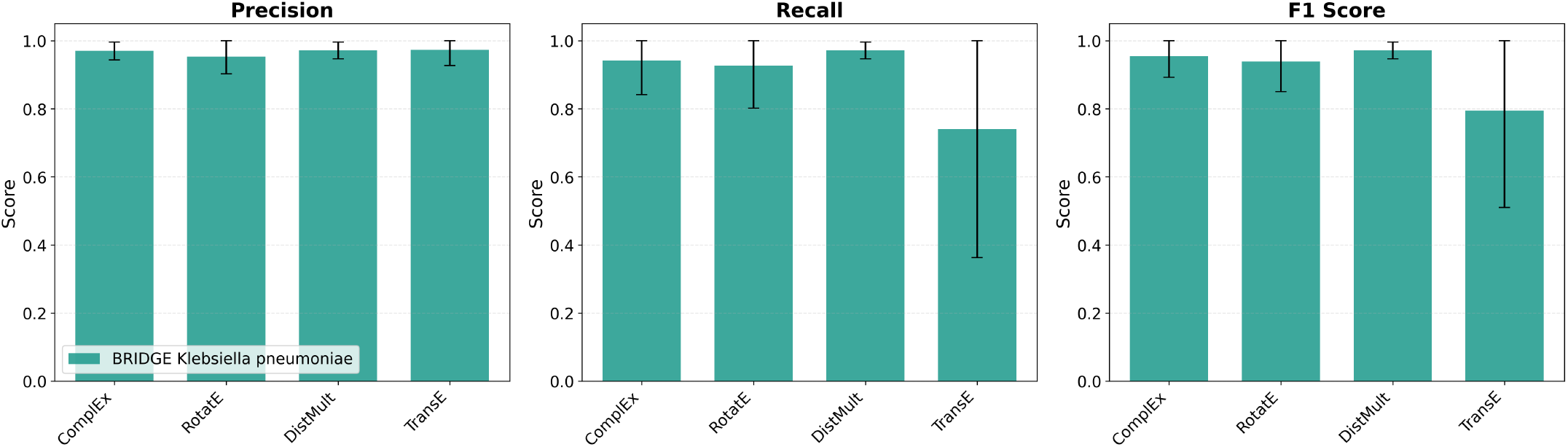
BRIDGE performance across different KGE models’ embeddings (as MLP inputs) for the *K. pneumoniae* KG.

**Fig. 6.**
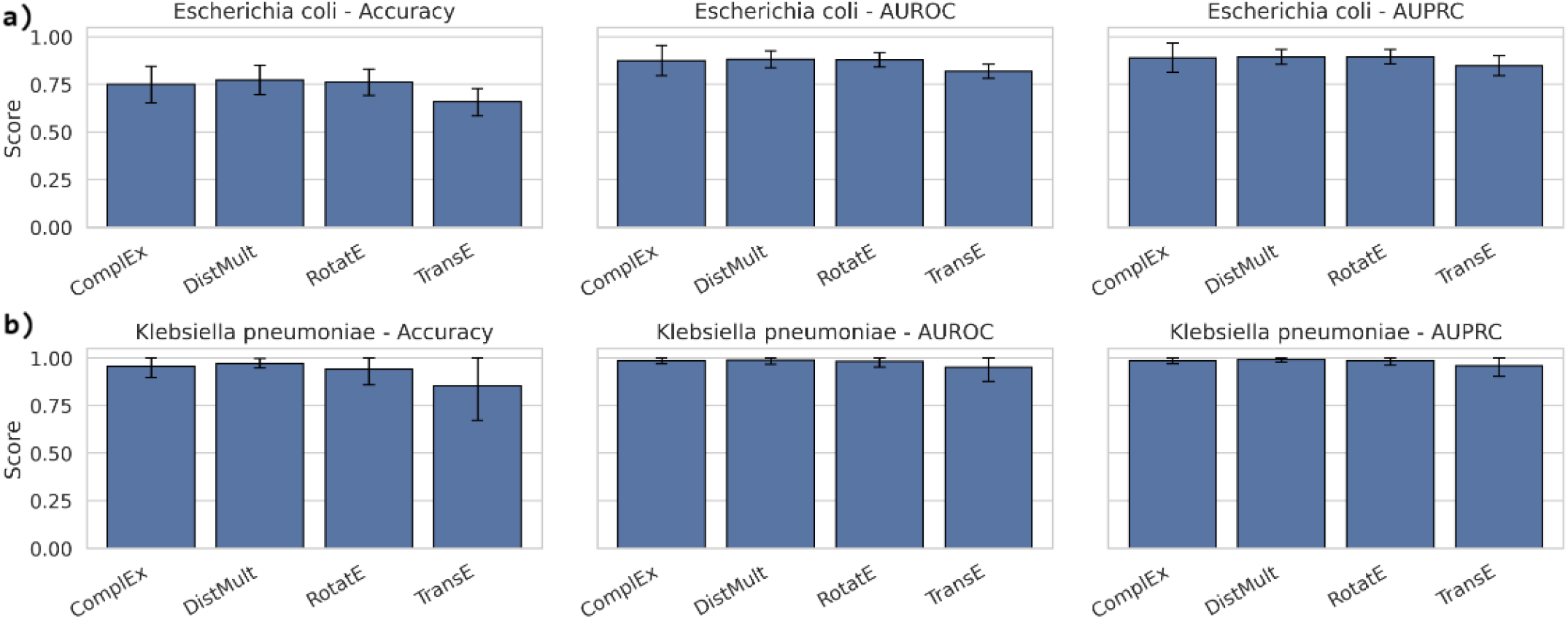
BRIDGE classification performance: a) accuracy, AUROC, and AUPRC scores for *E. coli* KG, b) accuracy, AUROC, and AUPRC scores for *K. pneumoniae* KG.

### E. Novel links

Tables IV and V summarise the verifiable positive and negative ARG-drug associations identified among the top 100 predictions generated by the ComplEx embedding model. These predicted links fall outside the scope of CARD, which serves as the sole source of curated ARG-drug relationships in the BRIDGE KG, and therefore represent previously unobserved candidate associations. To assess BRIDGE’s inductive capability, the model was provided with the complete KG and tasked with ranking novel ARG–drug links based on predicted likelihood.

Through a targeted literature review, we identified 26 verifiable predictions for the *K. pneumoniae* KG and 25 verifiable predictions for the *E. coli* KG. Most of these associations correspond to ‘Positive Links’, where independent studies report evidence supporting resistance conferred by the gene to the associated drug. A smaller subset of predictions were validated as negative links, indicating cases where the literature explicitly reports that the gene does not confer resistance to the corresponding drug.

Examination of the predictions in Table IV suggests that the model tends to prioritise associations that align with well-characterised biological mechanisms. For example, several high-confidence predictions involve links between members of the KPC gene family and ceftazidime. While CARD documents this association for a limited subset of KPC variants, existing literature indicates that additional variants may exhibit similar resistance profiles. These observations suggest that the model may capture shared biological signals across related gene families. In contrast, comparable patterns are less evident among the predictions in Table V, potentially reflecting differences in biological complexity, data availability, or annotation coverage for *E. coli*-associated resistance mechanisms.

**TABLE IV.** Novel Predicted Links for the *K. pneumoniae* KG.

| **Gene** | **Drug** | **Link Type** | **Citation** |
|---|---|---|---|
| *K. pneumoniae* PBP3 mutants | ceftazidime | Positive Link | [43] |
| *K. pneumoniae* OmpK36 | cefoxitin | Positive Link | [44] |
| CMY-2 | imipenem | Positive Link | [45] |
| KPC-79 | ceftazidime | Positive Link | [46] |
| CMY-2 | cefepime | Negative Link | [47] |
| *K. pneumoniae* OmpK36 | ertapenem | Positive Link | [48] |
| KPC-71 | ceftazidime | Positive Link | [49] |
| KPC-36 | ceftazidime | Positive Link | [46] |
| KPC-72 | ceftazidime | Positive Link | [46] |
| VEB-25 | ceftazidime | Positive Link | [50] |
| KPC-74 | ceftazidime | Positive Link | [51] |
| *K. pneumoniae* OmpK36 | imipenem | Negative Link | [52] |
| KPC-52 | ceftazidime | Positive Link | [46] |
| KPC-54 | ceftazidime | Positive Link | [46] |
| KPC-95 | ampicillin | Positive Link | [46] |
| *K. pneumoniae* OmpK35 | cefotaxime | Positive Link | [53] |
| KPC-58 | ceftazidime | Positive Link | [46] |
| OXA-1 | ceftazidime | Positive Link | [54] |
| KPC-73 | ceftazidime | Positive Link | [46] |
| KPC-25 | ceftazidime | Positive Link | [46] |
| KPC-35 | ceftazidime | Positive Link | [46] |
| *K. pneumoniae* OmpK35 | meropenem | Positive Link | [55] |
| KPC-65 | ceftazidime | Positive Link | [46] |
| KPC-2 | ceftazidime | Positive Link | [56] |
| KPC-41 | ceftazidime | Positive Link | [46] |
| OKP-B-3 | ceftazidime | Negative Link | [57] |

**TABLE V.** Novel Predicted Links for the *E. coli* KG.

| **Gene** | **Drug** | **Link Type** | **Citation** |
|---|---|---|---|
| AAC(6')-Ib-cr6 | gentamicin | Negative Link | [58] |
| APH(3')-IIa | amikacin | Positive Link | [59] |
| TEM-30 | ceftazidime | Positive Link | [60] |
| APH(3')-IIa | tobramycin | Positive Link | [61] |
| CTX-M-27 | ceftriaxone | Positive Link | [62] |
| rmtB | neomycin | Positive Link | [63] |
| CTX-M-15 | cefixime | Positive Link | [64] |
| H-NS | tigecycline | Negative Link | [65] |
| rmtE2 | neomycin | Positive Link | [66] |
| mphA | spiramycin | Positive Link | [67] |
| rmtB | streptomycin | Positive Link | [68] |
| marA | ceftazidime | Positive Link | [69] |
| AAC(6')-I33 | sisomicin | Positive Link | [70] |
| rmtE | neomycin | Positive Link | [66] |
| oqxB | tetracycline | Positive Link | [71] |
| npmA | spectinomycin | Positive Link | [72] |
| npmA | steptomycin | Negative Link | [72] |
| rmtE2 | tobramycin | Positive Link | [73] |
| rmtB | ribostamycin | Positive Link | [74] |
| oqxA | tetracycline | Positive Link | [75] |
| mdtM | ciprofloxacin | Positive Link | [76] |
| NDM-4 | ertapenem | Positive Link | [77] |
| mdtE | ciprofloxacin | Positive Link | [78] |
| AAC(3)-IId | streptomycin | Positive Link | [79] |
| acrD | novobiocin | Positive Link | [80] |

BRIDGE’s inductive potential is reflected in the fact that approximately 20% to 30% of the top-ranked predictions could be corroborated by existing literature, despite being absent from both the training data and CARD’s curated knowledge base. At the same time, a comprehensive quantitative assessment of BRIDGE’s inductive performance remains challenging, as many predicted associations are currently unverifiable due to limited published evidence and the extensive coverage of known AMR-drug relationships within CARD.

Taken together, these results provide preliminary qualitative support for BRIDGE’s ability to generate biologically plausible hypotheses beyond transductive link prediction. However, further experimental validation and longitudinal curation efforts will be necessary to rigorously assess false positive rates and fully characterise the framework’s inductive predictive capabilities.

### F. Ablation Study

To assess the extent to which the learnt embeddings benefit from specific relationship types, we conducted an ablation study. In this analysis, at most one relation type is removed from the KG at a time, allowing us to isolate and quantify the contribution of each relation to overall model performance. After removing each relation type in turn, the KGE models were retrained and evaluated using standard metrics, including MRR, MR, and Hits@k. The results of this experiment were validated using five-fold cross-validation.

As shown in Figure 7, relations such as *is part of family* and *interacts with* had a strong influence on model performance, with their removal causing up to a 28% drop in MRR in some models. Removal of relationships like *is equivalent to* or *confers resistance to drug class* slightly improved results. No single relationship’s removal improved performance consistently across bacteria or model, encouraging its inclusion in the KG. While this is not an exhaustive study, it is indicative that the relationships included in the KG are certainly not redundant and contribute (at least in certain cases) to forming a more accurate latent space of embeddings.

**Fig. 7.**
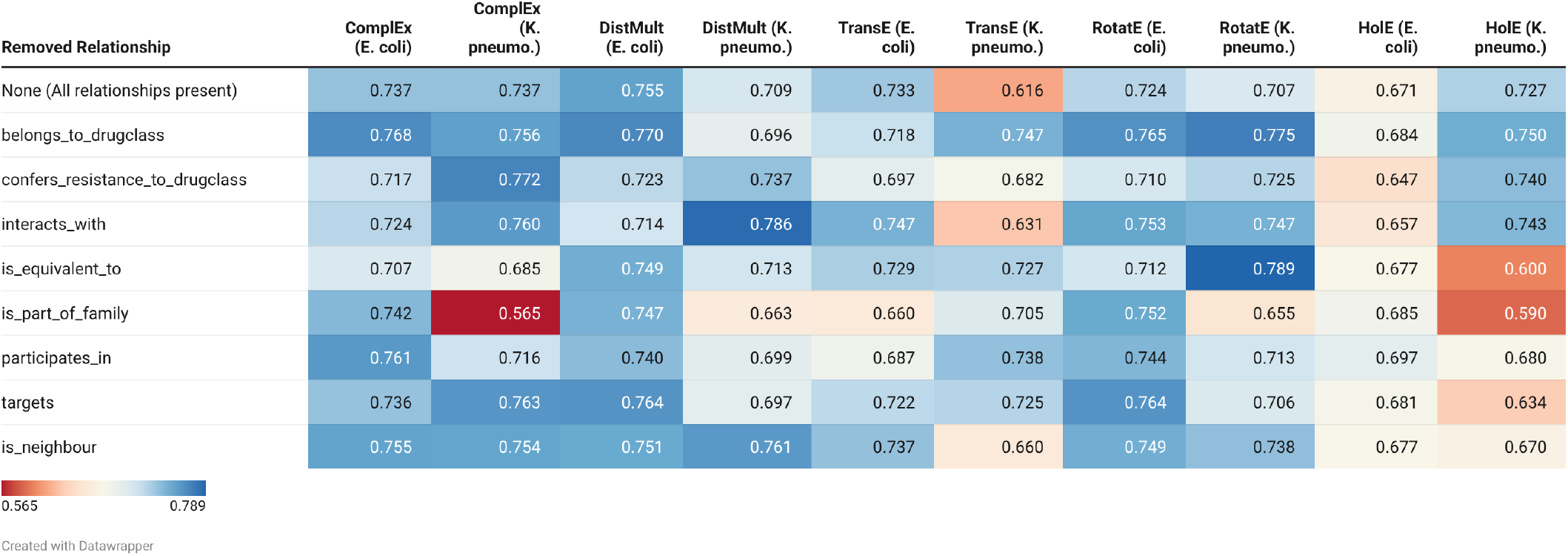
MRR scores for E. coli and K. pneumoniae across removed relationships and different models.

## V. Discussion

This study shows that curating a biologically grounded KG for the prediction of antimicrobial resistance that integrates gene neighbourhood, PPIs and drug-target relations, among other relationships, makes link prediction highly effective, particularly when using semantic-matching embedding models appended with an MLP classifier. Furthermore, across both *K. pneumoniae* and *E. coli*, semantic models outperformed distance-based models on MRR and Hits@k. The curation of the KGs for both bacterial species has been a controlled and measured affair as shown in Figure 1.

Figures 4 and 5 depict the range of precision, recall and F1 scores achieved by the BRIDGE framework and its clear improvement over KIDS. An important consideration is that the evaluation datasets differ, as the reported KIDS performance is taken directly from their original study. This difference reflects the fact that both KIDS and BRIDGE are end-to-end frameworks in which data curation is an integral component. While KIDS constructs a larger and more comprehensive KG by integrating a broader set of data sources, BRIDGE focuses on a more targeted graph enriched with PPIs and gene neighborhood information, enabling more context-aware modeling despite a smaller graph size.

While curated KGs minimise inconsistencies, the method has a tendency to limit the size of the resulting KG. Relying on thresholds for STRING confidence and BLASTp identity can restrict interactions included in the KG. Ultimately, the problem at hand is broken down into an embedding problem, followed by an MLP. The embeddings alone yield scalar scores indicating triplet plausibility upon their combination to form the objective function, but they lack the flexibility to model non-linear decision boundaries or to output explicit class probabilities. The two-stage architecture balances the interpretability of KGEs with the predictive power of an MLP, which pure embedding scores may miss.

## VI. Conclusion and Future Work

Our proposed framework, BRIDGE, introduces a novel approach to constructing an AMR KG to uncover new AMR links through KGE and link prediction. We emphasise the meticulous construction of the KG to avoid the need for ineffective inconsistency resolution techniques. To take a step back, we used a KG because AMR is a multi-relational problem; it does not just matter whether two items are close in a distance-based model sense, but how they are connected through specific biological relations (gene neighbourhood, drug target, etc.). Encoding these relation types using KGE models gives BRIDGE meaningful context and improves link prediction when compared to models that rely solely on a single feature space. In addition to incorporating conventional AMR KG relations and nodes, our approach integrates gene neighbourhood information annotated by BAKTA, as well as drug-target interactions and PPIs. This inclusion of biologically relevant information is expected to enhance the predictive capabilities of our framework. The KGEs were then concatenated and passed through a neural network classifier, allowing the model to capture more complex patterns and to convert a ranking problem into a classification task. The results show a clear improvement over the baseline KIDS framework. The BRIDGE framework outperforms baseline methods in every metric, with the DistMult-BRIDGE model reaching a highest mean accuracy of 77.27% and AUPRC of 89.41% for *E. coli* and an accuracy of 97.13% and AUPRC of 99.07% for *K. pneumoniae*, emphasising the need for careful data curation before KG prediction tasks.

Several promising directions remain for extending the BRIDGE framework. First, while this study focuses on *K. pneumoniae* and *E. coli*, future work will aim to scale the framework to additional bacterial species and strains, enabling broader evaluation of generalisability across diverse genomic and resistance landscapes. Second, incorporating additional data modalities, such as gene and protein sequence embeddings derived from foundation models (e.g., BioBERT or protein language models), minimum inhibitory concentration (MIC) measurements, and temporal or environmental meta-data, could further enhance predictive power and biological interpretability. Third, more advanced negative sampling strategies and uncertainty-aware learning could be explored to better distinguish hard negatives and reduce false positives, particularly in inductive prediction settings. Finally, experimental validation of high-confidence novel predictions, in collaboration with wet-lab or clinical partners, will be essential to rigorously assess real-world applicability and establish BRIDGE as a decision-support tool for antimicrobial resistance surveillance and drug discovery.

## Acknowledgment

We thank Dr Robert G. Beiko for his constructive feed-back and helpful suggestions during the preparation of this manuscript. A.K. was supported by the Mitacs Globalink Research Internship, and Y.K. was supported by the Saint Mary’s Dean of Science Summer Award. This research was enabled in part by support provided by ACENET (www.ace-net.ca) and the Digital Research Alliance of Canada (www.alliancecan.ca). The work reported in this study has been partially supported by an NSERC (Natural Sciences and Engineering Research Council) Discovery Grant (RGPIN-2020-05869).

---

**Aravind Iyer** is currently pursuing his Masters Degree (Sc.M) in Computer Science from Brown University and is expected to graduate in 2025. He has been a research assistant to Dr. Rita-mbhara Singh in 2025. He has worked as a research intern for Dr. Somayeh Kafaie’s Lab at Saint Mary’s Univeristy, Halifax, Canada. His research interests include genomics, deep learning, knowledge graphs, and bioinformatics. Mr. Krishnan was a recipient of the Mitacs Globalink Research Internship Award in 2024 and is also a co-recipient of the IEEE Research Alchemist award from the IEEE Bangalore chapter.

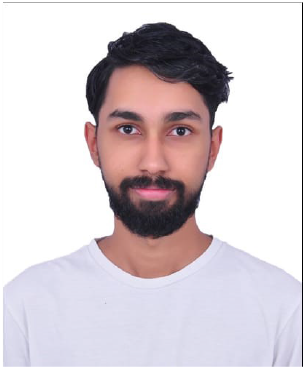

**Yusuff Kazeem** is a graduate of Computer Science and Mathematics from Saint Mary’s University, Halifax, Canada. His research interests include knowledge graphs, machine learning, bioinformatics, and mechanistic interpretability, with a particular focus on the intersection of data-driven methods and healthcare applications.

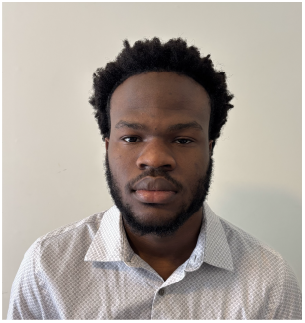

**Enayat Rajabi** earned his M.Sc. degree in Soft-ware Engineering from Ferdowsi University of Mashhad, Iran, in 2004. He then completed his Ph.D. (Cum Laude) in Information and Knowledge Engineering at the University of Alcalá, Spain, in 2015. From 2015 to 2017, he was a Postdoctoral Fellow at Dalhousie University in Halifax, Canada, where he conducted research on semantic web analytics frameworks for healthcare systems. After that, he worked as a Data Scientist at REDspace in Halifax, where he developed several data platforms utilizing artificial intelligence and machine learning. Dr. Rajabi has published numerous articles in computer science journals, focusing primarily on AI, machine learning, and knowledge graphs. Currently, he is an Associate Professor of Business Analytics at the Shannon School of Business, Cape Breton University, Nova Scotia, Canada. He has secured research grants from the Natural Sciences and Engineering Research Council of Canada (NSERC) and Research Nova Scotia. Additionally, he serves as a principal and co-principal investigator on various AI projects in healthcare and has received several “Excellent” ratings for his academic theses.

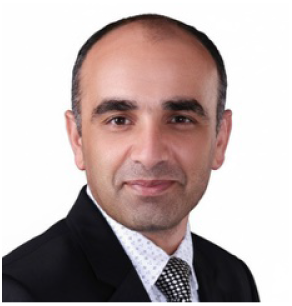

**Somayeh Kafaie** received the M.Sc. degree in Computer Engineering (Software) from Iran University of Science and Technology, Iran (2011), and the Ph.D. degree in Computer Engineering from Memorial University of Newfoundland, Canada (2017). She was a postdoctoral fellow in the Department of Computer Science at Memorial University (2017–2019) and in the Faculty of Computer Science at Dalhousie University (2020–2021). She has been an Assistant Professor in the Department of Mathematics and Computing Science at Saint Mary’s University, Halifax, Canada, since 2019. Her research interests include complex networks, knowledge graphs, explainable AI, and bioinformatics. She has received several awards, including the Best Editor Award from IEEE Wireless Communications Letters, the Fellow of SGS distinction, and the GSU Award for Excellence in Teaching.

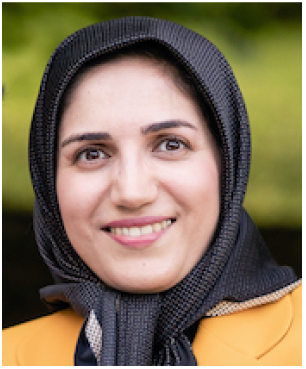

